# Polyanionic Non-Collagenous Proteins and Their Analogues Promote Artificial Mineralization of Embryonic Mouse Bone

**DOI:** 10.1101/2025.06.10.658745

**Authors:** Muhammad Wisnugroho, Fraser H. J. Laidlaw, Andrei V. Gromov, Colin Farquharson, Fabio Nudelman

**Author notes:** Department of Physics, The University of Indonesia, Biophysics Laboratory, Depok, Indonesia (Muhammad Wisnugroho). **Corresponding Author Fabio Nudelman –** School of Chemistry, The University of Edinburgh, Joseph Black Building, Edinburgh, UK.

## Abstract

Non-collagenous proteins (NCPs) are specialized biomacromolecules within the extracellular matrix (ECM) that regulate the mineralization of calcified tissues, such as bone and dentin. Numerous *in vitro* studies have demonstrated that natural polyanionic NCPs and their analogues can mediate intrafibrillar mineralization, characterized by the infiltration of apatite minerals into collagen fibrils. However, these studies primarily utilize self-assembled collagen fibrils or demineralized mature tissues, leaving it unclear whether pristine embryonic bone ECM at a developmental stage permissive to mineral deposition can regulate intrafibrillar mineralization independently or requires polyanionic NCP substitutes to promote the process artificially. To address this, we employed an *ex vivo* model of endochondral ossification using metatarsals isolated from 15-day-old embryonic mice (E15). In addition to a supersaturated calcium (Ca) and inorganic phosphate (Pi) medium, we introduced fetuin-A, a native polyanionic NCP or poly-DL-aspartic acid (pAsp), commonly used as an NCP substitute. The incorporation of either additive was essential for the effective mineralization of embryonic metatarsals. Both fetuin-A and pAsp played a direct role in facilitating the infiltration of Ca-Pi precursors into the avascular cartilaginous matrix. Raman spectroscopy and electron microscopy confirmed the formation of hydroxyapatite (HAp) exhibiting diverse levels of crystallinity, with fetuin-A supplementation resulting in the greatest HAp accumulation within the rudiments. HAp was localized in the perichondrium, a region conducive to initial mineralization and enriched with a fibrillar network of collagen types I and II. Three-dimensional reconstructions implementing Dijkstra’s algorithm revealed the association between HAp and collagen fibrils either organized in an intrafibrillar, extrafibrillar, or combined arrangement.

## Introduction

Bone is a calcified connective tissue that stores minerals, creates the skeletal system, and actively contributes to metabolic functions.^1^ In mammals, all long bones are formed via endochondral ossification, which is a replacement of hyaline cartilage anlagen with bone tissue.^2^ Collagen fibrils—the primary structural units of cartilage and bone—are internally strengthened by minerals at the nano-structural level via a process called mineralization to improve their structural integrity and resistance to internal and external loads.^3^ Bone minerals consist of platelet-shaped hydroxyapatite (HAp) crystals with a length, width, and thickness of 12–100 nm, 10–40 nm, and 0.61–5 nm, respectively.^4–7^ However, biogenic HAp typically exhibits a non-stoichiometric phase, characterized by a high carbonate (CO_3_^2–^) ion content, ranging from approximately 5.8% to 7.4% by weight, along with minor substitutions of other anions and cations, such as Na^+^, Mg^2+^, K^+^, Cl^−^, and F^−^.^8,9^ This imbalance makes biological HAp highly reactive and effective at exchanging calcium (Ca), inorganic phosphate (Pi), and other ions, as bones are intricately connected with blood vessels and constantly exposed to blood flow.^10^

The complex “intrafibrillar mineralization” mechanism by which nano-sized HAp crystals nucleate and grow within collagen fibrils during bone tissue formation has been studied for decades.^3,11–13^ Recent *in vitro* experiments have successfully achieved intrafibrillar mineralization by treating reconstituted collagen fibrils with supersaturated Ca and Pi solution with poly-DL-aspartic acid (pAsp) as a negatively charged “polyanionic” non-collagenous proteins (NCPs) substitute.^14,15^ This polyanionic macromolecule was demonstrated *in vitro* to interact with Ca and Pi ions in solution and prevent spontaneous nucleation of HAp crystals by stabilizing amorphous calcium phosphate (ACP) precursor phase via a polymer-induced liquid precursor (PILP) process.^14,16,17^ This negatively charged precursor interacts with a positively charged region at the C-terminus end of the gap zones of collagen fibrils, which facilitates the infiltration of ACP within the fibril and subsequent ACP to HAp transformation.^18^ The PILP process was further demonstrated to be used as a biomimetic system to study different parameters controlling the mineralization process, such as the role of the polymer molecular weight^19^ and extended beyond single fibril to promote the mineralization of dense collagen scaffolds and the remineralization of demineralized bone.^20,21^

Advanced remineralization studies using decalcified and chemically fixed mature dentin tissue have shown that the native extracellular matrix (ECM), which includes collagen fibrils and other associated macromolecules contains the information needed to guide mineralization when exposed to metastable Ca-Pi solution, resulting in the formation of ACP within the tissue.^22^ These experiments highlight the need to determine whether the ECM of developing bone possesses inherent functional properties that regulate mineralization or whether the presence of polyanionic NCPs is necessary to trigger mineral deposition within collagen fibrils.

Fetuin-A is an embryonic NCP with polyanionic characteristics, classified as a phosphorylated glycoprotein (42–68 kDa) and a member of the cystatin superfamily.^23–25^ It consists of two structurally linked cystatin D1 and D2 domains, along with a third domain rich in proline and glycine.^26^ At the onset of endochondral ossification, hypertrophic chondrocytes and cartilage ECM exhibit high expression of fetuin-A, suggesting its potential involvement in tissue mineralization.^25,27^ Additionally, this polyanionic NCP normally circulates in the blood and extracellular fluid with a half-life of several days, to regulate the Ca-Pi precursors and prevent spontaneous vascular calcification.^28,29^

Previous comparative *in vitro* studies demonstrated that fetuin-A promotes intrafibrillar formation of HAp crystals in both the reconstituted collagen fibrils and decalcified bone tissue via mineralization by an inhibitor size exclusion (ISE) mechanism.^12^ During the ISE process, fetuin-A binds selectively to Ca-Pi precursors^30^ and inhibits the apatite precipitation outside the collagen fibrils.^12^ This allows the precursors to exclusively infiltrate the fibril and subsequently nucleate to form HAp crystals within the intrafibrillar compartments.^12^ The ISE route contrasts with the PILP process in which a liquid ACP precursor phase forms extrafibrillarly and enters the collagen fibrils via capillary forces.^14^ Despite the differences, both mechanisms highlight the intricate interplay between collagen fibrils, polyanionic NCP substitutes, and Ca-Pi minerals to control and facilitate mineralization.^12,14^

Nevertheless, all those studies were conducted either using self-assembly collagen fibrils^13–15^ or decalcified mature tissue.^12,22^ Hence, it remains unclear whether the pristine and unmineralized ECM of embryonic bone contains sufficient capability to regulate the mineralization directly or whether native polyanionic NCPs and similarly charged macromolecule analogues are necessary to promote artificial mineralization within embryonic bone. Accordingly, our present study aimed to investigate these issues using an *ex vivo* model of endochondral ossification in embryonic mouse metatarsals, an organ explant culture pioneered by Burger and colleagues^31^ with a novel upgrade on the incorporation of natural polyanionic NCP or its polymer analogue. We hypothesized that the polyanionic-type macromolecules: fetuin-A and pAsp are crucial to initiate mineral precursors infiltration and promote artificial mineralization within embryonic bone tissue. It is important to understand the role of these negatively charged macromolecules and identify the best promoter of collagen mineralization during embryonic bone formation to aid the development of novel bio-inspired materials that are greatly beneficial in promoting fracture repair and treating bone-related diseases.

## Results

### Development of Embryonic Bone Mineralization Model

The 15^th^ gestational days (E15) metatarsals were isolated as previously described.^32^ At this developmental stage, no matrix mineralization was observed at mid-diaphysis (**Figure 1A**). Moreover, the rudiments are still avascular and translucent, with proliferating/hypertrophic chondrocytes, collagen type-II fibrils, and aggrecan-type proteoglycans as their primary composition.^33,34^ While metatarsals at E14 consist almost entirely of small-sized chondrocytes, the widening of lacunae or the beginning of cartilage hypertrophy is seen in the mid-diaphysis of E15, which is the preliminary sign that the tissue is ready to accept mineral deposition.^31^ The cartilage matrix is pristine and unmineralized, and the bone collar is still absent compared to later development stages.^31,32^ Small amounts of soft tissue, such as tendons and muscles remain attached to the outside of the rudiments, which are very difficult to remove during dissection. However, these soft tissues remove themselves after a few days in culture.

**Figure 1.**
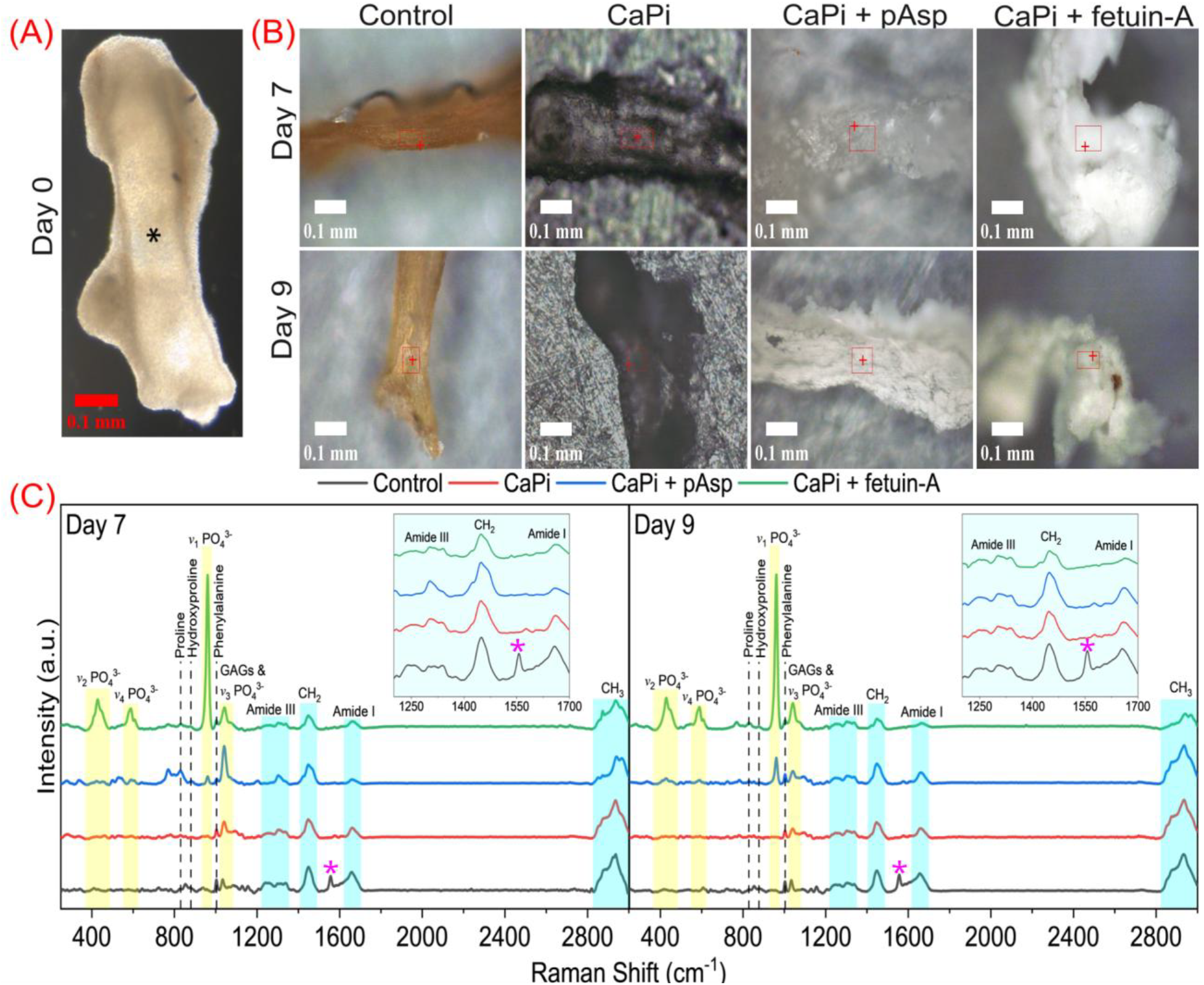
Optical images of E15 metatarsals on (A) day 0 in which the mineralized core was absent at the mid-diaphysis (black asterisk) and (B) after 7 and 9 days-of-culture. The red boxes indicate the Raman mapping area in each sample. (C) Raman spectra of E15 metatarsals cultured for 7 and 9 days with each line representing a different treatment: Control (black), CaPi (red), CaPi + pAsp (blue), and CaPi + fetuin-A (green) (Inset: amide bands of collagen in 1200–1700 cm^−1^ range). Amide II band (purple asterisk) was only visible in the Control samples at both incubation days.

From this point onward, artificial mineralization of E15 metatarsals was conducted using four different incubation mediums at two observation times: 7 and 9 days-of-culture. The culture mediums were classified as cell culture medium in the absence of CaPi minerals (Control), medium with CaPi minerals only (CaPi), and medium with CaPi minerals plus polyanionic additives (CaPi + pAsp and CaPi + fetuin-A).

### Physical Appearance Change and Mineralization Features

After 7 and 9 days in culture, the E15 metatarsals treated with CaPi + pAsp and CaPi + fetuin-A became stiff and appeared white under the microscope compared to the Control and CaPi samples (**Figure 1B**). Since all samples were washed thoroughly with water before observation, this physical transformation may indicate mineral deposition within the metatarsals. It should be noted that the metatarsals cultured in the CaPi medium (i.e., supersaturated Ca and Pi solution with respect to HAp stoichiometry^35,36^) did not appear white under dark field (DF) optical imaging, which reflect the absence of mineral within the tissue. Comparatively, there were fewer solid precipitates observed in the bottom of the culture well of both samples supplemented with pAsp and fetuin-A than in the CaPi-only sample. Altogether, this white color transformation, an increase in rudiment stiffness, and less mineral sedimentation in the culture well are the first indication of a mineralization within E15 metatarsals treated with NCP substitutes.

### Organic and Inorganic Phases Identification

To investigate both the organic and inorganic phases of E15 metatarsals, Raman observation on a selected region of the mid-diaphysis (**Figure 1B**) was conducted after 7 and 9 days-of-culture. The resulting 100 spectra from each sample mapping area were averaged and normalized with respect to amide III band (~1250 cm^−1^) in agreement with previous studies.^37^ The combined Raman spectra in the range of 250–3000 cm^−1^ region was displayed for both culture days (**Figure 1C**). The vibrational band assignments were identified and described (Supporting Information: **Table S1**).

The peaks corresponding to the collagen, namely amides, along with the ring of proline, hydroxyproline, and phenylalanine were present in all samples both at 7 and 9 treatment days (**Figure 1C** and Supporting Information: **Table S1**). Amide I and III bands are indicative of the collagen phase signatures andwere observed across all treatment conditions. In contrast, amide II band, which is associated with other proteins and lipids was exclusively detected in the Control sample on both incubation days. The presence of glycosaminoglycans (GAGs) could also be identified by the peaks corresponding to the pyranose ring at 1045 cm^−1^ and the symmetry O–SO_3_^−^ bond at 1062 cm^−1^. Importantly, the peaks correspond to the *v*_1_, *v*_2_, *v*_3_, and *v*_4_ modes of phosphate (PO_4_^3−^) were present only in the samples incubated in CaPi + fetuin-A with little change from 7 to 9 days-of-culture. The appearance of these peaks indicates the presence of HAp crystals within the metatarsals. The spectrum of samples incubated in CaPi + pAsp after 7 days only displayed a low-intensity peak at 961 cm^−1^, corresponding to the *v*_1_ mode of PO_4_^3−^, whose intensity increased slightly after 9 days. In addition, the *v*_2_ and *v*_4_ peaks were absent at 7 days and at low intensity at 9 days of incubation in CaPi + pAsp, indicating that mineralization was slower than with CaPi + fetuin-A, with less HAp formed. The absence of the *v*_1_ and *v*_4_ modes of PO_4_^3−^ in the Control and CaPi samples showed that without additives no mineralization of the embryonic bone tissue takes place.

Taken together, these results imply that fetuin-A and pAsp can both promote artificial mineralization of E15 metatarsals, with fetuin-A being more effective. It should be highlighted that Raman spectroscopy is a surface technique, with a beam penetration depth of approximately up to 12 µm. Therefore, electron microscopy (EM) examination is necessary to evaluate mineral localization deep inside the metatarsals (discussed below).

### Coils Arrangement and Intrafibrillar Mineralization

A more ordered coils structural arrangement of collagen fibril is correlated to intrafibrillar mineralization, or it can also be interpreted as fibril preparation to receive minerals deposit.^38^ The quantification of coil arrangement level (CAL) values (**Equation 1**) on Raman spectra resulted in the highest overall value on CaPi + fetuin-A treatment on both 7 and 9 days of culture. The CAL mean values (Supporting Information: **Table S2**) were increased as follows: Control (0.511 ± 0.08) < CaPi + pAsp (0.667 ± 0.07) < CaPi (0.669 ± 0.13) < CaPi + fetuin-A (1.078 ± 0.06) for day 7, and Control (0.534 ± 0.09) < CaPi (0.641 ± 0.13) < CaPi + pAsp (0.705 ± 0.13) < CaPi + fetuin-A (1.411 ± 0.16) for day 9. With prolonged incubation time, the CAL variability of the additive-supplemented samples increased, signifying that the fibrils became more ordered with time in culture as a result of mineral deposition within the fibrillar compartment.

Statistical analysis revealed that samples with CaPi only and CaPi plus additives have a significant increase in CAL (*p* < 0.001) compared to the Control sample (**Figure 2A**), and a significant increase (*p* < 0.001) in the value for CaPi + fetuin-A from day 7 to day 9 of culture (**Figure 2B**). The absence of PO ^3−^ peaks in the CaPi sample suggests that, even though no mineral was formed, enough Ca and Pi infiltrated into the tissue to induce coil arrangement changes in the collagen fibrils. These observations also demonstrate the direct role of fetuin-A and pAsp in facilitating the infiltration of Ca-Pi precursors into the avascular cartilage rudiments to affect coil arrangements and promote mineralization.

**Figure 2.**
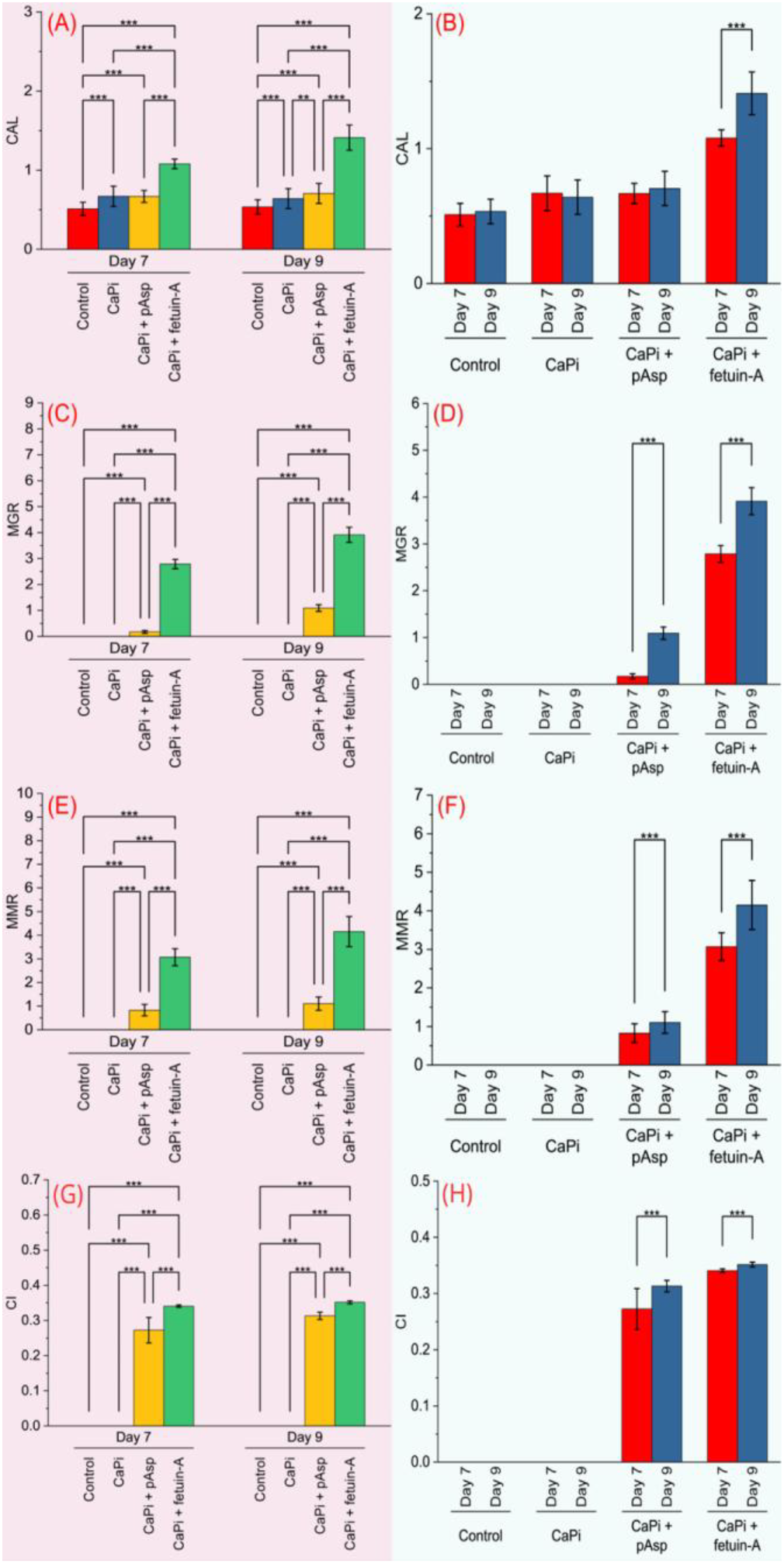
Statistical analysis of E15 metatarsals culture with (A–B) CAL, (C–D) MGR, (E–F) MMR, and (G–H) CI as compared to each specific medium (left side) and culture time (right side). Zero (null) values for MGR, MMR, and CI in the Control and CaPi samples due to the absence of *v*_1_PO_4_^3−^ peaks. \**p* < 0.05; \*\**p* < 0.01; \*\*\**p* < 0.001.

### Mineral Development in Relation to Tissue Mineralization

To investigate the correlation between the formation of HAp crystals and the collagen matrix, Raman quantitative analyses on several aspects (**Figure 2C–H**) were determined and compared. Tissue mineralization was evaluated by both the mineral to GAGs ratio (MGR) and the mineral to matrix ratio (MMR) values. The MGR distribution within the mineralized matrix of the GAGs-rich ECM (**Equation 2**) depicted that while all samples with NCP substitutes have mineral deposition in their ECM, there were differences in the mineralization level between them; with fetuin-A supplemented samples producing the highest overall MGR value on both culture days. The MGR mean values (Supporting Information: **Table S2**) were increased as follows: Control = CaPi (null) < CaPi + pAsp (0.171 ± 0.05) < CaPi + fetuin-A (2.785 ± 0.18) for day 7, and Control = CaPi (null) < CaPi + pAsp (1.091 ± 0.13) < CaPi + fetuin-A (3.912 ± 0.29) for day 9. All additive-treated samples were significantly different (*p* < 0.001) from Control and CaPi samples (**Figure 2C**). Moreover, the MGR of the pAsp and fetuin-A supplemented samples increased with time in culture (*p* < 0.001) suggesting a greater mineral deposition into the tissue with prolonged culture time (**Figure 2D**).

Furthermore, when the overall quantity of minerals was compared to the total collagen matrix (MMR, **Equation 3**), the CaPi + fetuin-A sample on both culture days also yielded the highest overall value when compared with the other treatment conditions. The MMR mean values (Supporting Information: **Table S2**) were increased as follows: Control = CaPi (null) < CaPi + pAsp (0.827 ± 0.24) < CaPi + fetuin-A (3.071 ± 0.36) for day 7, and Control = CaPi (null) < CaPi + pAsp (1.104 ± 0.28) < CaPi + fetuin-A (4.151 ± 0.64) for day 9. The MMR for the CaPi + fetuin-A sample was higher (*p* < 0.001) when compared to the CaPi + pAsp sample (**Figure 2E**). This also implies that the rate of mineralization is faster in the presence of fetuin-A than pAsp. Accordingly, when the culture time was prolonged to 9 days, the ratio increased (*p* < 0.001) further for both fetuin-A and pAsp treated samples (**Figure 2F**), indicating the increase of tissue mineralization level during this period.

Mature minerals have higher crystallinity values due to the crystal apatite having a more perfect lattice organization with fewer ionic substitutions or stoichiometric arrangement.^39^ The crystallinity index (CI) (**Equation 4**) of the CaPi + fetuin-A samples was greater than that of the other samples at both culture time points. The CI mean values (Supporting Information: **Table S2**) were increased as follows: Control = CaPi (null) < CaPi + pAsp (0.271 ± 0.04) < CaPi + fetuin-A (0.341 ± 0.01) for day 7, and Control = CaPi (null) < CaPi + pAsp (0.313 ± 0.01) < CaPi + fetuin-A (0.352 ± 0.01) for day 9. The lower CI in the pAsp compared to the fetuin-A supplemented samples (*p* < 0.001) is reflective of a well-defined and organized HAp crystal structures with fetuin-A addition (**Figure 2G**). The respective CI of both NCP supplemented samples increased (*p* < 0.001) from day 7 to day 9 of the culture suggesting that crystal growth and maturation was a time dependent process (**Figure 2H**).

In general, tissue mineralization (MGR and MMR) has a parallel correlation to mineral crystallinity (CI). All quantitative analyses yielded the same results, in which the CaPi + fetuin-A sample at both culture days resulted in the highest overall value in contrast to the other treatments. Concisely, artificial HAp crystals formation inside E15 metatarsals was successfully initiated and promoted by adding natural polyanionic NCP, such as fetuin-A (1 mg/mL) or its polymer substitute, such as pAsp (25 µg/mL) in a supersaturated Ca (2.5 mM) and Pi (1 mM) solution. Collagen-associated mineralization occurred in fetuin-A supplemented cartilage as indicated by the highest coil structural change and tissue mineralization is correlated with mineral crystallinity, which reveals the interplay between collagen fibrils, polyanionic NCP substitutes, and Ca-Pi minerals.

### Mineral Localization in Cartilaginous Tissue

Transmission electron microscopy (TEM), scanning-transmission electron microscopy (STEM), and energy dispersive X-ray spectroscopy (EDX) observations were used to investigate mineral localization in the tissue sections. In accordance with the Raman measurement, mineral presence was absent in the Control and CaPi samples after 7 and 9 days of culture (**Figure 3– TEM)**. Mineral was sparsely detected in the metatarsals in CaPi + pAsp sample at 7 and 9 days-of-culture. In contrast, CaPi + fetuin-A revealed the presence of minerals throughout the inner region (i.e., near cartilage core) of the metatarsals. These TEM findings have also confirmed that mineralization takes place inside the metatarsals rather than just on their surface. Minerals in the CaPi + pAsp and CaPi + fetuin-A samples have similar plate-shaped crystals with a width/thickness of about 5–20 nm and a length of 30–100 nm. The minerals of the CaPi + fetuin-A samples were more neatly distributed along the collagen fibrillar network, a pattern not so obvious in the CaPi + pAsp samples. In agreement with the previous data, this indicates that fetuin-A is a robust promoter of crystal apatite deposition.

**Figure 3.**
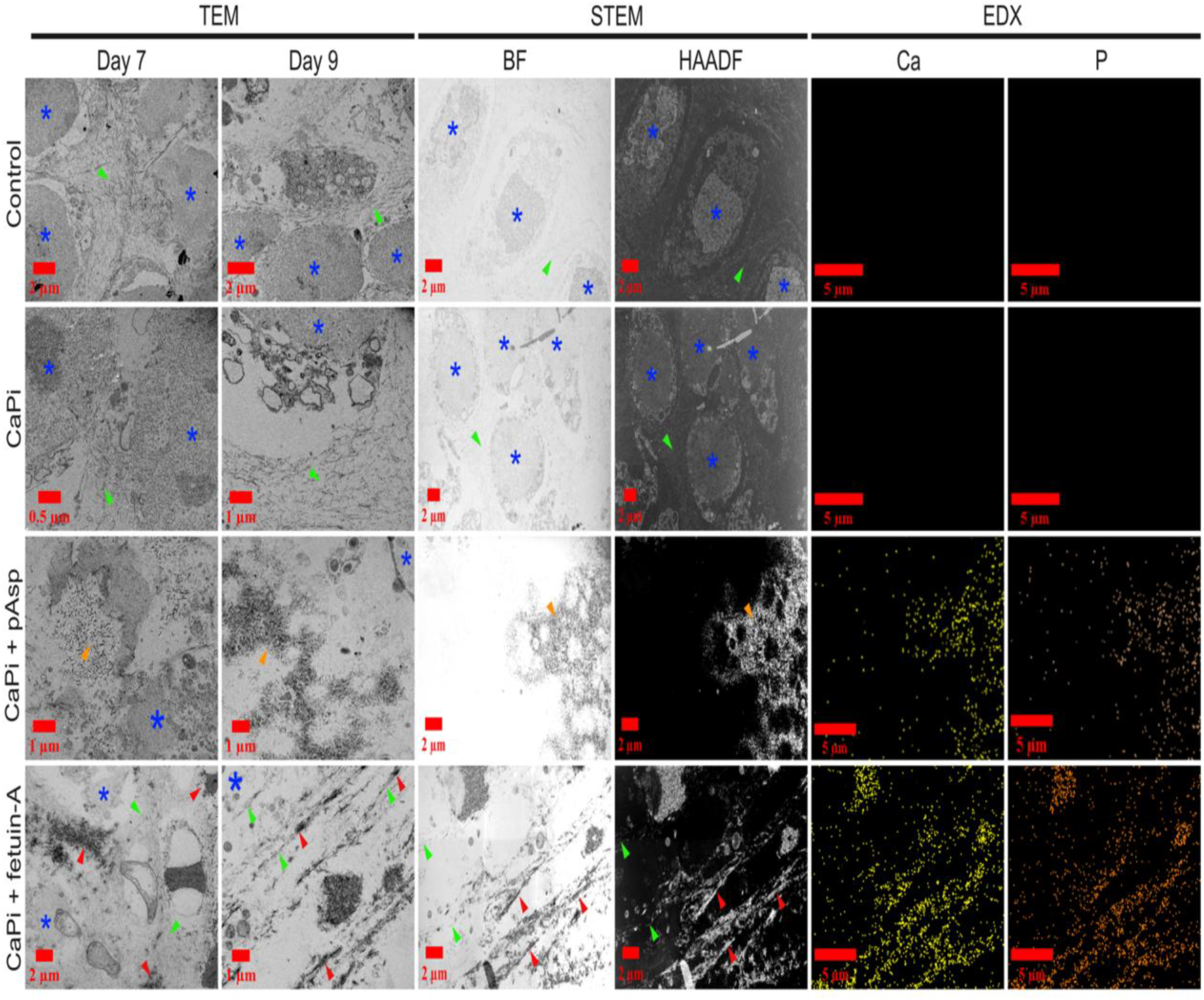
Comprehensive TEM (day 7 and day 9) and STEM-EDX (day 9) observations on the inner region (i.e., near cartilage core) of E15 metatarsals cultures. Irregular-shaped cells (blue asterisks) were adjacent to the unmineralized collagen fibrillar (green arrowheads) network in Control and CaPi samples. The presence of minerals (orange arrowheads) not associated with collagen fibrils was detected in the CaPi + pAsp samples. A distinguishable pattern between unmineralized (green arrowheads) and mineralized (red arrowheads) collagen fibrils was observed in CaPi + fetuin-A supplemented samples, which matched with their Ca and P distribution. BF: bright field; HAADF: high angle annular dark field; STEM: scanning-transmission electron microscopy; EDX: energy dispersive x-ray.

Elemental mapping after 9 days-of-culture of the samples with NCP additives showed the presence of both Ca and P, confirming that the tissue is deposited with calcium-phosphate mineral (**Figure 3–EDX**). Combined with bright field (BF) and high angle annular dark field (HAADF) imaging, the analysis indicated that in the CaPi + fetuin-A samples, the mineral was arranged adjacent to collagen fibrils (**Figure 3–STEM**). Overall, the mineral within the CaPi + fetuin-A sample was shown to localize to the perichondrium, forming an electron-dense layer about 30–40 µm thick (**Figure 4A**). As the perichondrium in E15 metatarsals contains both collagen types-I and -II fibrils, the mineralized collagen type-I fibrils within this layer represents the initiation of the bone collar, which fully forms at later stages of bone development.^31,34,40,41^ Within CaPi + fetuin-A supplemented metatarsals, some regions of the collagen fibril (*d*~20 nm) were mineralized, and a cloud-like mineral complex was attached to the fibril surface (**Figure 4B**). These findings are in agreement with the role of fetuin-A in promoting collagen mineralization.^12,18^

**Figure 4.**
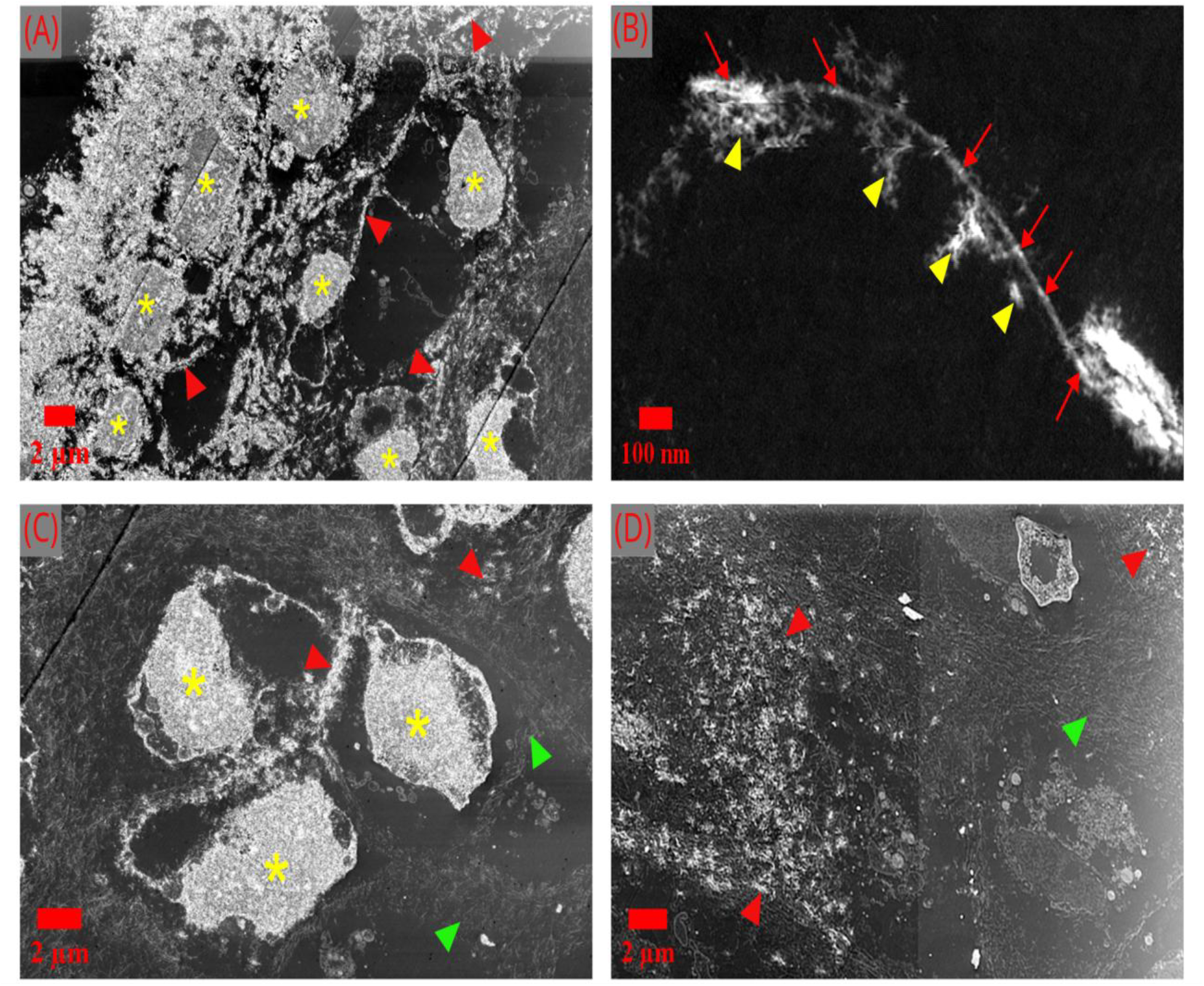
Selected STEM (HAADF) images of E15 metatarsals culture with CaPi + fetuin-A addition. (A) The highly mineralized perichondrium layer extends to the inner region of the cartilage tissue, where numerous calcified cells (yellow asterisks) reside. (B) The partially mineralized collagen fibril (red arrows) with the mineral-fetuin-A complex adhered to the surface in a cloud-like structure (yellow arrowheads). At the cartilage core, minerals were also heavily localized in (C) the cells and (D) some regions of the collagen matrix either still unmineralized (green arrowheads) or mineralized (red arrowheads).

Interestingly, not only the collagen matrix but also the cells within the cartilage core of fetuin-A treated samples were heavily calcified. The cells were relatively small (*d*~4–8 µm) with irregularly shaped morphologies (**Figure 4C–D**). These cells possibly undergo shrinkage, necrosis, and apoptosis because the culture conditions were insufficient to maintain cell survival, which was followed by rapid mineral deposition and HAp crystallization within their apoptotic cytoplasm.

### Three-Dimensional (3D) Structure of Mineralized Embryonic Metatarsal

To reconstruct and visualize the calcified collagen matrix, focused-ion beam scanning electron microscopy (FIB-SEM) tomography was conducted on CaPi + fetuin-A treated samples since these were the only ones that contained overt areas of mineralization. Large area mapping of a cross section through the metatarsal using the backscattered electron (BSE) revealed the distribution of HAp (**Figure 5A**), and two areas of the perichondrium at the mid-diaphysis were selected as the region of interests (ROIs) (**Figure 5A inset**) for tomography. In contrast to the initial E15 metatarsal (**Figure 1A**) before culture (*L*~0.85–1 mm), the cartilage rudiments become relatively longer (*L*~1.2– 1.5 mm) after being cultured with CaPi + fetuin-A (**Figure 5A**), indicating tissue growth. However, it is important to note that this growth is minimal (Δ*L*~0.2–0.5 mm) in comparison to the typical elongation of cultured embryonic cartilage rudiments during endochondral ossification (Δ*L*~1.5–2 mm), likely due to culture conditions that were insufficient to support prolonged cell survival.^31,32^

**Figure 5.**
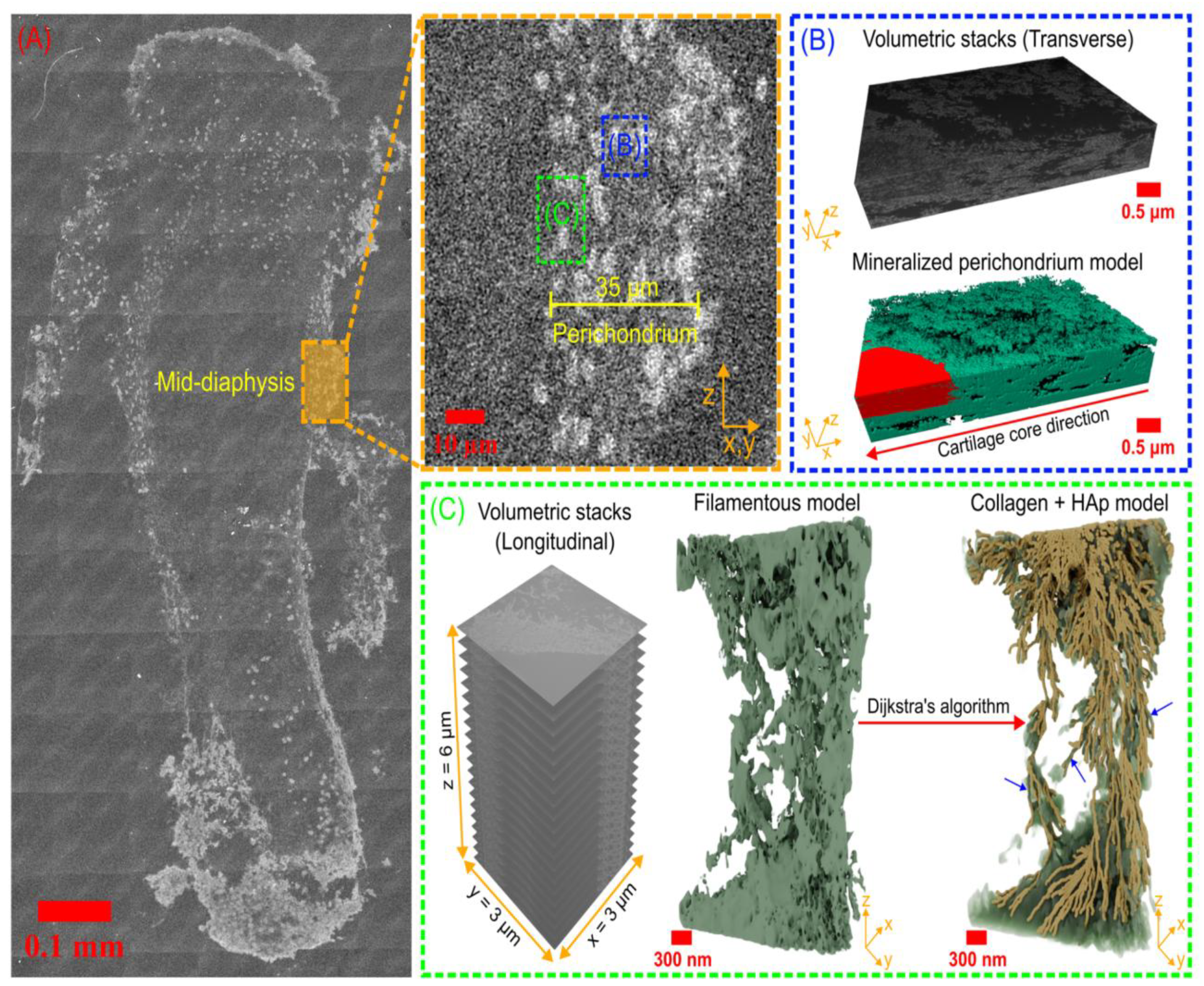
(A) Overlay area of E15 metatarsals cultured with CaPi + fetuin-A. Within the perichondrium of the mid-diaphysis (inset: close up area), the ROIs were selected for serial surface imaging in (B) transverse (blue dashed box) and (C) longitudinal (green dashed box) orientations (*x, y*, and *z* axes match between the direction in the inset and each 3D model). The representative 3D models from volumetric stacks were displayed as (B) a mineralized perichondrium model composed of the irregular cell (red) embedded within mineralized ECM (green) and (C) a filamentous model composed of calcified collagen fibrils orientated parallel to the perichondrium. The shortest path method or Dijkstra’s algorithm implementation on the filamentous model exposes a structure of collagen fibrils (yellow) embedded within minerals (green semi-transparent volume). Intrafibrillar HAp crystals deposited within the projected collagen fibrillar network are indicated by blue arrows.

A 3D reconstruction of the transverse volumetric stacks showed that the perichondrium layer was heavily mineralized in the lateral direction towards the cartilage core at the mid-diaphysis (**Figure 5B**). A cytoplasm portion of the irregular-shaped cell adjacent to the perichondrium layer was fully embedded in mineralized ECM, which is portrayed as a 3D mineralized perichondrium model. Parallel to the perichondrium layer a filamentous structure was observed from longitudinal image stacks and displayed as a 3D filamentous model (**Figure 5C**). This filamentous structure was likely constructed from numerous collagen fibrils that aggregated into a collagen fiber together with HAp crystal deposited intra- or extrafibrillarly. It was difficult to differentiate between collagen fibrils/fibers and HAp crystals as they were closely packed together with similar image confluency and contrast. However, implementing a Dijkstra’s—shortest path method—algorithm^42^ (Supporting Information: **Figure S2**) to follow the trace or common path of the collagen fibrils and combining it with the previous model, it was revealed that numerous smaller filamentous-like structures and rough apatite morphology were responsible for the 3D collagen + HAp model at the perichondrium (**Figure 5C** and Supporting Information: **Video S1**). Since collagen fibrils often interweave, branch, and overlap, this algorithm purposely computes optimal paths within two or more adjacent voxels that minimize intensity differences or follow geometric constraints through noisy BSE data. When the structural paths are discontinuous and the volumetric morphology is spherical at specific site, the algorithm is unable to reconstruct fibrils, resulting in HAp classification. Therefore, each filament structure was identified as a collagen fibril that had a length of approximately 1–6 µm. All these collagen fibrils were interconnected and embedded within dense HAp crystals.

According to this 3D model, these HAp crystals were portrayed to be associated with collagen fibrils either organized in an intrafibrillar, extrafibrillar, or combined arrangement. This model projection also reinforces the TEM and STEM results (**Figure 3** and **Figure 4**), indicating that HAp crystals in CaPi + fetuin-A samples were organized in a regular pattern along collagen fibrillar network. It should be highlighted that extrafibrillar crystal arrangement is predominant within the perichondrium at this time studied. Nevertheless, further evaluations are necessary to determine the exact location of HAp crystal deposition.

## Discussion

Our *ex vivo* embryonic metatarsal culture provides a highly suitable and physiological ECM for investigating early bone mineralization. This model more accurately replicates the initial bone tissue mineralization process compared to earlier *in vitro* approaches, such as the mineralization of self-assembled collagen fibrils or the remineralization of demineralized mature bone tissue. Uncalcified embryonic metatarsal tissue at E15 provides an authentic and functional cartilaginous ECM template for mineralization.^31,32^ Moreover, the absence of blood vessels in the tissue at this gestational age infers that the mineral infiltration solely depends on a simple diffusion or convection mechanisms.^33,34,40,43^ These characteristics create an excellent model for investigating the functional role of embryonic bone ECM and polyanionic NCP substitutes in regulating Ca-Pi precursors to promote tissue mineralization during endochondral ossification. To the best of our knowledge, this is the first *ex vivo* attempt to utilize natural polyanionic NCP or its analogue to promote artificial mineralization of the embryonic long bone.

During embryonic bone development, the blood carries the Ca-Pi ions within the vascular network in its supersaturated and balanced state with respect to HAp stoichiometry, [Ca] = 2.05– 2.58 mM and [Pi] = 0.78–1.49 mM.^35,36^ The avascular cartilage anlagen takes up the mineral precursors from the surrounding extravascular fluid after they are released from adjacent capillaries via paracellular pathways^44,45^ and larger negatively charged macromolecules (42–68 kDa), such as albumin and fetuin-A enter the interstitial fluid through the opening of the transendothelial channel to continuously maintain the Ca-Pi ionic equilibrium and prevent spontaneous apatite nucleation.^46–48^ Accordingly, we chose two polyanionic macromolecules known to regulate the mineral precursors *in vitro*: pAsp (25 µg/mL with approximate size of 11 kDa or *d*~2.9 nm) and fetuin-A (1 mg/mL with approximate size of 64 kDa or *d*~5.3 nm) in addition to the serum-equivalent Ca and Pi concentrations of 2.5 mM and 1 mM, respectively.^12,14,35^

The first intriguing point of discussion is the inability of E15 cartilage rudiments to mineralize spontaneously upon exposure to a supersaturated CaPi solution, despite the rudiments possessing a functional ECM and being at a developmental stage conducive to mineral deposition. It must be noted that these conditions are different from those of normal culture conditions (e.g. tissue grown in 95% air and 5% CO_2_ supply^31,32^) where matrix mineralization continues for several days. In this latter case, the cells play a critical role in controlling matrix homeostasis and mineral deposition. In the case of our system, mineralization only occurred within the tissue in the presence of either fetuin-A or pAsp along a supersaturated CaPi medium. The hydrated state of cartilage maintains its structural integrity by intermolecular electrostatic-steric forces predominantly between negatively charged chondroitin sulfate networks within the GAGs chain.^49^ Together with small cartilage pores (*d*~6–14 nm), these properties may provide high resistance to fluid flow and water redistribution on the tissue because of low permeability.^50–52^ Moreover, in unloaded cultures, the mineral infiltration rate into the tissue is also impaired in compliance with previous studies.^53^ Therefore, it is unlikely that supersaturated Ca-Pi medium alone can promote spontaneous cartilage mineralization as observed in our work. Without a mineral regulator, like fetuin-A or pAsp, Ca-Pi precursors nucleate faster and spontaneously precipitate within the solution rather than diffuse into the cartilage anlagen.

There is a possibility that when the Ca and Pi concentrations fall below those selected in our study (i.e., [Ca] < 2.5 mM and [Pi] < 1 mM), the precursors may not undergo spontaneous nucleation in solution, allowing them to infiltrate the tissue without the necessity of either fetuin-A or pAsp as a mineralization regulator. However, a cartilage ECM has been shown to tolerate substantially higher Ca concentrations in the presence of normal Pi levels without inducing apatite precipitation, a phenomenon not observed in aqueous solutions.^54,55^ Under these conditions, precursors would merely diffuse in and out of the tissue, making mineralization unfeasible.

Native polyanionic NCPs and their analogues, such as fetuin-A and pAsp inhibit crystal nucleation and precipitation in solution, subsequently promoting intrafibrillar mineralization of collagen *in vitro*.^12,14^ Our results show that the incorporation of fetuin-A or pAsp within Ca and Pi-rich fluid is necessary to initiate mineral precursor infiltration and promote embryonic bone mineralization. One possible explanation is that these polyanionic macromolecules may form a semi-liquid or amorphous complex with the Ca-Pi precursors in solution as widely postulated.^12,14^ This amorphous “ACP-NCPs” complex may adhere to the cartilage rudiment surface and progressively diffuse through the pores of a dense perichondrium layer to promote mineralization.

Our Raman measurements at the mid-diaphysis region disclosed that the embryonic cartilage supplemented with additives beside CaPi solution, especially with fetuin-A treatment resulted in similar spectra (**Figure 1C**) to the bone, even when compared to calcified cartilage.^56,57^ Additionally, significant changes to the coil structural arrangement (**Figure 2A–B**) indicated that the collagen fibrils become more ordered due to the deposition of crystals within the fibrils thus aligning with previous studies.^38^ It is noteworthy that CaPi-only treatment also yielded a change in the coil structure without mineralization and the value is comparable to that observed in the CaPi + pAsp treated rudiments. In contrast, the CaPi + fetuin-A gave a higher value that may account for the association of the mineral with collagen fibrils, implying this association was also absent in the CaPi + pAsp samples. Although electron microscopy in combination with 3D reconstruction showed the mineral following a filamentous structure, indicative of its association with collagen fibrils (**Figure 5C**), it could not allow for a clear distinction of whether it is deposited within the gap zones or on the fibril surface. Thus, we concur that the HAp crystals either organized in an intrafibrillar, extrafibrillar, or combined arrangement.

It is possible that in the presence of CaPi only, small quantities of mineral precursors can enter the cartilage rudiments and alter the coil arrangement in the ECM. However, native NCPs, such as chondrocalcin, osteopontin, osteocalcin, and bone sialoprotein are possibly absent in the cartilage tissue at this stage of development (i.e., primary ossification) and only synthesized at a later ossification stage in accordance to previous *in vivo* studies.^58–61^ Therefore, it is likely that the unexpected mineral precursors that bypass the tissue hydraulic permeability barrier and infiltrate the cartilage anlagen without the support of polyanionic additives cannot crystallize properly within collagen fibrils or they are only sufficient to alter the coil structural order moderately.

We reveal that the perichondrium of the developing “cartilage to bone” anlagen is the specific region that is heavily mineralized (**Figure 4A**). Our finding is similar to *in vivo* observations that endochondral ossification of long bone begins with the formation of a mineralized “bone collar” layer around this site, which contains overlapping collagen type-I and type-II composition at this gestational period.^31,34,40^ This finding is not surprising because it is established that type-II collagen plays an important role during endochondral ossification of bone tissues.^62^ Recent *in vivo* observations on the endochondral ossification of mice auditory ossicles and the bony labyrinth tissues demonstrated that mineralization with high mineral density take place within the conjoint type-I and -II collagen network, suggesting a potential link between the presence of collagen type-II and hypermineralization of bone during endochondral ossification.^63^ Proteoglycans associated with collagen type-II exhibit a higher molecular mass compared to those bound to collagen type-I fibrils.^64^ Given the propensity of Ca ions to localize at proteoglycan sites,^65,66^ the presence of collagen type-II within the osteoid matrix—predominantly consisting of collagen typeI fibrils—may facilitate enhanced mineral deposition. This notion is supported by our observation of a highly mineralized region within the perichondrium, consistent with the elevated mineralization typically associated with type-I and -II collagen-containing tissues.

An important aspect to consider is which type of collagen in the cartilage anlagen contains the intrafibrillar deposition of crystal apatite. Although collagen type-II has a similar 67 nm periodic spacing as collagen type-I in accordance to previous studies,^67^ collagen type-II cannot be mineralized intrafibrillarly *in vitro* by using pAsp.^68^ Moreover, *in vivo* cartilage mineralization studies have also demonstrated that early crystal nucleation is localized within extrafibrillar sites, where proteoglycans are heavily distributed and not within collagen type-II fibrils^.65,69^ Therefore, we propose that if intrafibrillar mineral deposition occurs inside E15 cartilage rudiments as the collagen becomes more ordered following the increase of CAL values and as illustrated by the 3D model within perichondrium, it likely occurs exclusively within collagen type-I fibrils.

Our work reveals that fetuin-A promotes embryonic bone mineralization more effectively than pAsp. Despite the similar polyanionic characteristics of pAsp to the acidic domain of NCP, this macromolecule is not an NCP like fetuin-A. We assume that fetuin-A or other natural NCPs have specialized mechanisms for regulating the mineral precursors in the tissue, which are beyond the electrostatic attraction between mineral ions and the charged surface of macromolecules.^70^ Furthermore, we considered the possibility that both pAsp and fetuin-A facilitate collagen mineralization of embryonic bone through the PILP process.^14^ However, the Raman spectra obtained are distinctly different between each additive with the PO_4_^3−^ intensity of pAsp significantly lower compared to fetuin-A treatment. Smaller pAsp molecules complexed with Ca-Pi precursors would be expected to penetrate faster and deeper into the dense-packed collagen fibrils. If the PILP mechanism is the optimal way for embryonic bone mineralization, pAsp should have a higher promoting influence than fetuin-A and not otherwise. Hence, we highlight that the ISE process, which emphasizes the regulation of mineralization dynamics as the most favourable route during embryonic bone mineralization, and this aligns with previous studies.^12^

Overall, we demonstrate that artificial mineralization of embryonic cartilage anlagen can be initiated and promoted with an external mineral supply and polyanionic NCP substitutes. Tissue mineralization is concentrated within the perichondrium, and the HAp crystals are associated with collagen fibrils either organized in an intrafibrillar, extrafibrillar, or combined arrangement. Nevertheless, both intra-and extrafibrillar mineralization are fundamental to the development of embryonic bones and the determination of their final mechanical properties.

## Conclusions

This study represents the first demonstration of embryonic bone artificial mineralization by utilizing natural polyanionic non-collagenous protein or its analogue within an *ex vivo* model of endochondral ossification. While embryonic bone tissue possesses an inherent molecular composition and functional advantages, its extracellular matrix alone is insufficient to directly promote mineralization. However, the presence of a polyanionic macromolecule, such as fetuin-A or poly-DL-aspartic acid within a supersaturated calcium-phosphate medium exerts a significant and direct influence on promoting embryonic bone mineralization, potentially by stabilizing the amorphous mineral phase. The mineralized embryonic metatarsals have organic and inorganic phases that are comparable to the bone tissue with hydroxyapatite as the sole mineral phase according to Raman spectroscopy analysis. Electron microscopy observation combined with a three-dimensional reconstruction depicted that hydroxyapatite crystals are heavily localized within the perichondrium in association with collagen fibrils either organized in an intrafibrillar, extrafibrillar, or combined arrangement. Overall, fetuin-A is an effective promoter of early mineralization during embryonic endochondral ossification.

## Experimental Section

### Reagents and Solutions

All chemicals were purchased from Sigma-Aldrich (Dorset, UK), unless otherwise stated.

### Animal Welfare

Pregnant female C57BL/6 mice were purchased from Charles Rivers Lab (Kent, UK) and maintained under conventional housing conditions with 12 hours light/dark cycle. All animal experiments were approved by The Roslin Institute’s Animal Users Committee and the animals were maintained in accordance with UK Home Office guidelines for the care and use of laboratory animals.

### Isolation and Culture of Embryonic Mouse Metatarsals

Pregnant female mice were sacrificed by cervical dislocation and their E15 embryos were collected following decapitation in accordance with home office guidelines in the UK. The middle three metatarsals were isolated and dissected under a dissecting microscope in accordance with previous protocols.^32^ Throughout the dissection procedure, the metatarsals were kept under preparation medium, which composed of 0.8 mL alpha minimum essential medium (*α*-MEM) (without ribonucleosides), 10.45 mL sterile phosphate-buffered saline (PBS), and 22.5 mg bovine serum albumin (BSA) (Fraction V).

For the mineralization process, metatarsals were cultured in 24-well plates with each well containing one metatarsal in 600 µL of specific culture medium with the final concentration as described (Supporting Information: **Table S3**). Each specific culture medium was buffered to normal blood pH 7.4 with 0.25 mM hydroxyethylpiperazine ethane sulfonic acid (HEPES) solution and incubated in closed environment under physiological body temperature (37 °C) without a carbon dioxide (CO_2_) supply. The culture medium was changed every 3 days throughout the culture period and the rudiments were collected after 7 and 9 days of culture. Metatarsals were individually washed several times with ultrapure water to remove excess minerals on the surface and fixed with 2.5% glutaraldehyde in 0.1 M sodium cacodylate buffer solution pH 7.4 at 37 °C for 1 hour. Metatarsals were then stored in 0.1 M sodium cacodylate buffer solution pH 7.4 at 4 °C until required for processing.

### Raman Spectroscopy

Metatarsals were characterized using a Thermo Scientific DXR3 Raman microscope equipped with 785 nm laser using 10X objective with NA = 0.25 and 3 µm laser spot size. Each metatarsal was analyzed following optimal measurement parameters of 10 mW laser power and 12 s exposure time, with each spectrum as the average of 3 accumulations. Raman mapping of each metatarsal within an area of mid-diaphysis region was carried out using 5 µm step between measurement points with focus tracking under dark field (DF) imaging condition to produce a consisted minimum 100 spectra. Spectra were fluorescence corrected (5^th^ order) by the instrument OMNIC software to exclude background variations. A total of resultant 800 spectra (4 culture medium treatments for 2 time points) were then processed by OriginPro 2024b software for subsequent asymmetric least squares (ALS) background subtraction and 5^th^ order fast Fourier transform (FFT) filter smoothing.

For quantitative analysis, integrated peak areas (*∫*_*A*_) of the spectra were carefully selected (Supporting Information: **Figure S1** and **Table S4**) based on the Raman assessment studies of cartilage and bone tissue.^37,56,57,71^ The *∫*_*A*_ values under the normalized curve within each band region were determined by OriginPro 2024b software with interpolation to rectangle edges. Raman visualization was made by averaging and normalizing all representative spectra relative to amide III band intensity in accordance with previous studies.^37^

#### Coil arrangement level (CAL)

A relative quantification to measure structural changes in collagen fibril as a consequence of intrafibrillar mineralization.^38^ This value was calculated by dividing the *α*-helical (ordered) proteins band (Amide III) to the random-coil (disordered) proteins band (Amide I).

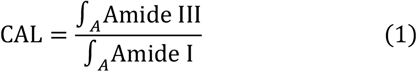

#### Mineral to GAGs ratio (MGR)

A relative estimation of the total mineralized matrix in comparison to ECM rich GAGs or relative amount of bone to cartilage tissue.^72,73^ This value was determined by dividing the pyranose ring-symmetric stretch of O-SO ^−^ band (GAGs) to the primary phosphate band (*v* PO_4_^3−^).

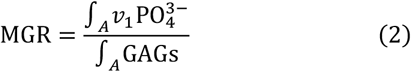

Since the GAGs band highly overlaps with the tertiary phosphate band (*v*_3_PO_4_^3−^) in agreement with previous studies^37^, all *∫*_*A*_GAGs values for each specific treatment were approximated from the subtraction with the *∫*_*A*_*v*_3_PO_4_^3−^ value of pure HAp mineral spectrum (rruff-database, https://rruff.info/hydroxylap-atite/R050512).

#### Mineral to matrix ratio (MMR)

A relative determination of the overall ratio between mineral and collagen in the tissue, which is correlated to the bone ash fraction.^74^ This value was calculated by dividing the primary phosphate band (*v*_1_PO_4_^3−^) to the *α*-helical (ordered) proteins band (Amide III).

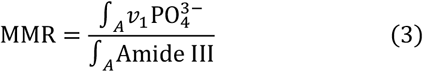

#### Crystallinity index (CI)

A relative quantification of apatite phase, which represents the average size of crystallites, perfection, and ordering in a crystal.^75^ This value was determined by using the width of the primary phosphate band, which is mathematically described as the reciprocal of the full width at half maximum (FWHM) of the *v*_1_PO_4_^3−^ band.

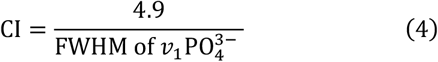

The value of 4.9 cm^−1^ corresponds to the average FWHM of the standard magmatic apatite (Conodont or phosphatic microfossil) and biological apatite have CI value in range of ≤ 0.5.^76^

#### Statistical analysis

A two-way analysis of variance (ANOVA) was performed with Bonferroni test to make a comparison of both culture treatments and culture times for each specific quantitative value. Differences were considered significant at *p* < 0.05, highly significant at *p* < 0.01, and very significant at *p* < 0.001.

### Electron Microscopy

For electron microscopy, metatarsals were post-fixed in 1% osmium tetroxide in 0.1 M sodium cacodylate buffer solution pH 7.4 for 45 minutes, then washed in three (10 minutes) changes of 0.1 M sodium cacodylate buffer solution pH 7.4. Next, specimens were dehydrated in 50%, 70%, 90%, and 100% ethanol three times for 15 minutes each, followed by two (10 minutes) changes in propylene oxide. Samples were then embedded in TAAB 812 epoxy resin.

#### Transmission electron microscopy (TEM)

The resin-embedded samples were cut (1 µm thickness) on a Leica Ultracut ultramicrotome, stained with toluidine blue, and viewed in a light microscope to select suitable areas for investigation. The ultrathin sections (60 nm thickness) were cut from selected areas and transferred to quantifoil TEM grids, stained in uranyl acetate and lead citrate solutions, then examined with JEOL JEM-1400 Plus TEM. Representative images were collected on a GATAN OneView camera.

#### Scanning-transmission electron microscopy (STEM) and energy dispersive X-ray spectroscopy (EDX)

Ultrathin sections (60 nm thickness) on quantifoil TEM grids were fitted on STEM grid holder, then viewed using Zeiss Crossbeam 550 equipped with a field-emission gun operation on dual channel STEM detector: bright field (BF) and high angle annular dark field (HAADF), operating at 20 kV accelerating voltage with 300 pA probes current and 3 mm working distance. Selected areas were further investigated for its elemental composition by EDX.

#### Focused-ion beam scanning electron microscopy (FIB-SEM) and serial surface imaging

The chosen sample in resin block was attached to the carbon-covered stub, mounted to the holder, then observed with Zeiss Crossbeam 550 field-emission gun (FEG) FIB-SEM using two detectors: secondary electron (SE) and backscattered electron (BSE), operating at 1 kV accelerating voltage with 200 pA probe current and 5 mm working distance. Large area mapping of the sample using the BSE was carried out first via ATLAS 5 software to produce an overlay area of the metatarsal and specific region of interest (ROI) at the perichondrium was selected. The stage was then tilted to 54^°^ to set the sample to be perpendicular to the FIB column. A platinum layer was deposited on top of ROI to protect the sample surface, then a trench was milled using FIB beam (30 kV; 15 nA) to create a cross-sectional of the ROI. After polishing of the ROI cross-section, automated serial milling and slice imaging took place to create both SE and BSE image stacks. Each slice (20 nm thickness) was generated with 300 pA probe current with scan speed = 4, N = 40, dwell time = 300 µs, and pixel size (X, Y) = 6.5 nm.

#### Image analysis and three-dimensional (3D) reconstruction

The obtained BSE image stacks were carefully aligned and segmented using Avizo 9.0 and Dragonfly software, respectively. The raw 3D model was then processed in Blender 3.6 software by a self-made geometry nodes (Supporting Information: **Figure S2**) based on Dijkstra’s shortest path algorithm.^42,77^ This algorithm was used to find the globally optimal fibril pathways between two or more selection (seed) points inside a given volume. The representative pathways were combined with the raw 3D model to create a new volumetric 3D model to illustrate and differentiate between inorganic minerals and collagen fibrils. The video of collagen fibrils pathway tracking and mineral localization (Supporting Information: **Video S1**) was generated and edited in Blender 3.6 software using the same image stacks segmentation of mineralized perichondrium layer in longitudinal direction which was transferred from Dragonfly software. The video was rendered using cycles render engine and OptiX denoising optimization with a high resolution of 1080p, 30 frames per second (fps), and very high contrast adjustment.

## Associated Content

### Data Availability Statement

The data that support the findings of this study are available from the authors upon reasonable request.

### Supporting Information

The Supporting Information is available.

Integrated peak areas (*∫*_*A*_) selection, Dijkstra’s shortest path geometry nodes, Raman spectra vibrational band assignments and quantitative assessment values, and final concentration of mineralization mediums (PDF).

Video S1 (MP4).

## Author Information

### Authors

**Fraser H. J. Laidlaw** – School of Physics and Astronomy, The University of Edinburgh, James Clerk Maxwell Building, Edinburgh, UK

**Andrei V. Gromov** – School of Chemistry, The University of Edinburgh, Joseph Black Building, Edinburgh, UK

**Colin Farquharson** – The Roslin Institute and Royal (Dick) School of Veterinary Studies, The University of Edinburgh, Easter Bush, Midlothian, UK

## Author Contributions

The manuscript was written through contributions of all authors. All authors have given approval to the final version of the manuscript.

## Acknowledgements

The authors acknowledge financial support from the Biotechnology and Biological Sciences Research Council (BBSRC) in the form of an Institute Strategic Programme Grant (BBS/E/RL/230001C). The authors thank the Indonesia Endowment Fund for Education (LPDP) from the Ministry of Finance, Republic of Indonesia for granting the scholarship and supporting this research. Electron microscopy data acquisitions were performed at the TEM (the Wellcome Trust Multi-User Equipment Grant WT104915MA) and Cryo FIB-SEM (EPSRC grant No. EP/P030564/1) facilities at the University of Edinburgh. The authors also thank Martin Singleton and Stephen Mitchell from Wellcome Trust Centre for Cell Biology, University of Edinburgh for their assistance regarding the TEM usage. For the purpose of open access, the authors have applied a CC-BY public copyright licence to any Author Accepted Manuscript version arising from this submission.

## Abbreviations

*α*-MEM: alpha minimum essential medium
Δ*L*: length difference
*∫*_*A*_: integrated peak areas
3D: three-dimensional
ACP: amorphous calcium phosphate
ALS: asymmetric least squares
ANOVA: analysis of variance
BF: bright field
BSA: bovine serum albumin
BSE: backscattered electron
Ca: calcium
CAL: coil arrangement level
CI: crystallinity index
CO_2_: carbon dioxide
CO_3_^2–^: carbonate
*d*: diameter
DF: dark field
E15: embryonic phase at 15^th^ days
ECM: extracellular matrix
EDX: energy dispersive X-ray spectroscopy
EM: electron microscopy
FEG: field-emission gun
FIB-SEM: focused-ion beam scanning electron microscopy
FFT: fast Fourier transform
FWHM: full width at half maximum
GAGs: glycosaminoglycans
HAADF: high angle annular dark field
HAp: hydroxyapatite
HEPES: hydroxyethylpiperazine ethane sulfonic acid
ISE: inhibitor size exclusion
*L*: length
MGR: mineral to GAGs ratio
MMR: mineral to matrix ratio
NCPs: non-collagenous proteins
pAsp: poly-DL-aspartic acid
PBS: phosphate-buffered saline
Pi: inorganic phosphate
PILP: polymer-induced liquid precursor
PO_4_^3−^: phosphate
ROIs: region of interest
TEM: transmission electron microscopy
SE: secondary electron
STEM: scanning-transmission electron microscop

