## Supporting Information Tables and Figures for "Polyanionic Non-Collagenous Proteins and Their Analogues Promote Artificial Mineralization of Embryonic Mouse Bone"

bioRxiv

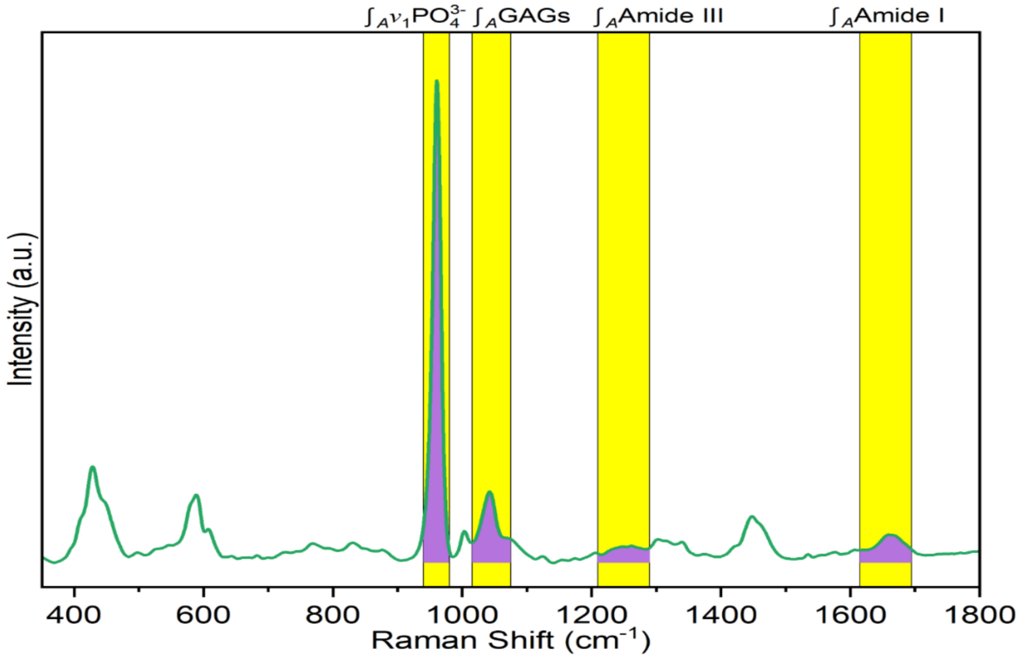


**Figure S1.** Integrated peak areas ($\int_A$) selection. Each $\int_A$ value (i.e., purple region under the Raman spectrum) represents the vibrational band of organic matrix and inorganic minerals in the embryonic tissue. Quantification details of $\int_A$ values are included in **Table S4**.

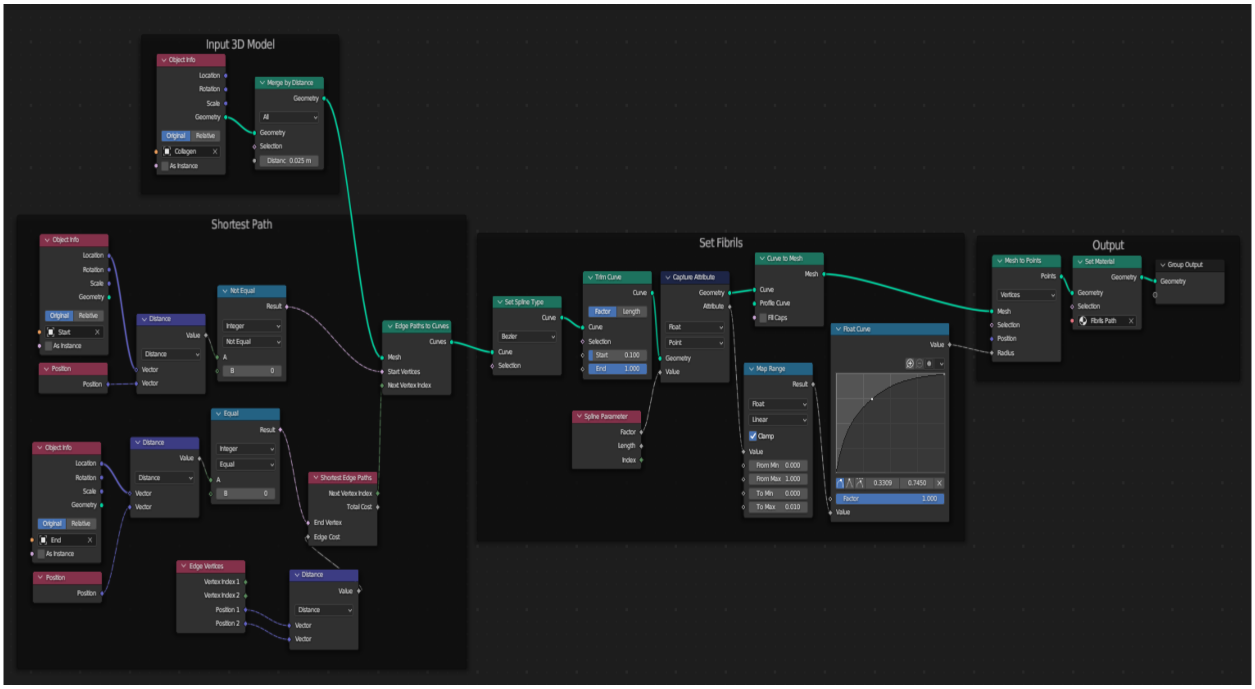


**Figure S2.** Shortest path geometry nodes. The implemented nodes are based on Dijkstra's algorithm.[1,2]

**Table S1.** The vibrational band assignments of Raman spectra observed in E15 metatarsals culture in agreement with previous studies.[3–9]

| Peaks ($cm^{-1}$) | Assignments | Components |
|---|---|---|
| 426–455 | (O–P–O) symmetry bend of $v_2PO_4^{3-}$ | HAp |
| 582–612 | (O–P–O) asymmetry bend of $v_4PO_4^{3-}$ | HAp |
| 830–855 | (C–C) stretch of proline ring | Collagen |
| 870–880 | (C–C) stretch of hydroxyproline ring | Collagen |
| 920–930 | (C–C) stretch of proline ring | Collagen |
| 960–962 | (P–O) symmetry stretch of $v_1PO_4^{3-}$ | HAp |
| 1002–1004 | (C–C) symmetry stretch of phenylalanine ring | Proteins; Collagen |
| 1039–1045 | (C–O–C) stretch of pyranose ring; (P–O) asymmetry stretch of $v_3PO_4^{3-}$ | GAGs; HAp |
| 1060–1063 | ($O–SO_3^-$) symmetry stretch of sulfated GAGs | GAGs |
| 1245–1275 | (C–N) stretch of amide III ($\alpha$-helix and $\beta$-sheet) | Collagen |
| 1440–1455 | $CH_2$ wagging | Proteins; Lipids |
| 1556–1560 | Amide II* | Proteins; Lipids |
| 1650–1685 | (C=O) stretch of amide I ($\beta$-sheet and random coil) | Collagen |
| 2856–3002 | $CH_3$ deformation/stretching | Proteins; Lipids |

*Amide II peak only visible in the Control sample

**Table S2.** The quantitative assessment values of Raman spectra in E15 metatarsals culture. Mean ± standard deviation from normal distribution.

| Time | Assessment | Control | CaPi | CaPi + pAsp | CaPi + fetuin-A |
|---|---|---|---|---|---|
| Day 7 | CAL | 0.511 ± 0.08 | 0.669 ± 0.13 | 0.667 ± 0.07 | 1.078 ± 0.06 |
| | MGR | null* | null* | 0.171 ± 0.05 | 2.785 ± 0.18 |
| | MMR | null* | null* | 0.827 ± 0.24 | 3.071 ± 0.36 |
| | CI | null* | null* | 0.271 ± 0.04 | 0.341 ± 0.01 |
| Day 9 | CAL | 0.534 ± 0.09 | 0.641 ± 0.13 | 0.705 ± 0.13 | 1.411 ± 0.16 |
| | MGR | null* | null* | 1.091 ± 0.13 | 3.912 ± 0.29 |
| | MMR | null* | null* | 1.104 ± 0.28 | 4.151 ± 0.64 |
| | CI | null* | null* | 0.313 ± 0.01 | 0.352 ± 0.01 |

CAL: Coil Arrangement Level; MGR: Mineral to GAGs Ratio; MMR: Mineral to Matrix Ratio; CI: Crystallinity Index.
*Zero (null) values for the Control and CaPi samples due to the absent of $v_1PO_4^{3-}$ peaks.

**Table S3.** The final concentration of each specific culture medium for mineralization of E15 metatarsals.

| Culture Medium | Final Concentration (per 600 µL) |
|---|---|
| Control | $\alpha$-MEM (without ribonucleosides) |
| CaPi | 2.5 mM $CaCl_2$ and 1 mM $K_2HPO_4$ |
| CaPi + pAsp | 2.5 mM $CaCl_2$, 1 mM $K_2HPO_4$, and 25 µg/mL pAsp |
| CaPi + fetuin-A | 2.5 mM $CaCl_2$, 1 mM $K_2HPO_4$, and 1 mg/mL fetuin-A |

**Table S4**. Parameters to determine the $\int_A$ values for all Raman spectra.

| Bands | Peaks ($cm^{-1}$) | Width ($cm^{-1}$) |
|---|---|---|
| $v_1PO_4^{3-}$ | 960 | 40 |
| GAGs | 1045 | 60 |
| Amide III | 1250 | 80 |
| Amide I | 1655 | 80 |
